# Timing of circadian shifts in light/dark preference by migratory bats supports targeting of sunset for magnetic compass calibration

**DOI:** 10.1101/2025.10.02.680110

**Authors:** Oliver Lindecke, Daniels Valerts, Viesturs Vintulis, Will T Schneider

## Abstract

Bats are nocturnal mammals whose migrations depend on the precise timing of activities to exploit favourable conditions, such as tailwinds and high insect activity, while evading predators. Understanding how migratory bats use sensory cues within their roosts to time emergence requires knowledge of the interplay between circadian rhythms, spatial cognition, and seasonal demands. We examined light-dark choice in tree-dwelling Nathusius’ pipistrelles (*Pipistrellus nathusii*) over an 18-hour cycle using a Y-maze. Bats’ preference for the lit exit increased prior to sunset, then switched back to preferring darkness only near sunrise. This behaviour contrasts with that of previously studied cave-dwelling bats. Irrespective of their exit choice, bats used echolocation in all trials, consistent with a multimodal sensory strategy in which light conveys critical environmental information beyond the range of echolocation, but not replacing it. However, light governed the time-dependent shift in exit selection. The circadian shifts in light/dark preference are consistent with two hypotheses related to navigation mechanisms and stopover behaviour: Preference for light cues may facilitate compass calibration mechanisms, whereas the dawn preference for darkness likely reflects roost-searching and predator avoidance. These findings underscore the integral role of vision in bat navigation and highlight how circadian rhythms modulate photic responses in a migratory context. Such insights are essential for designing wildlife-friendly lighting and for interpreting future multi-sensory experiments, including those probing bat magnetoreception, where natural photic responses must be taken into account.

## INTRODUCTION

Many animals possess a wide array of sensory abilities that allow them to process environmental cues and adjust behavioural responses. Sometimes, however, cues may confer contrasting information, or may be missing entirely. In these situations, animals depend on cue hierarchies where sensory inputs are given priority over others when cues are ambiguous (Ben-Ari and Inbar, 2014; Chase, 1983; Lebhardt et al., 2012; Moore and Phillips, 1988; Rossel et al., 1978; Stephenson, 2016), competing (Chase, 1981; Knight et al., 2014; Wiltschko et al., 1998a), or unavailable (Hagstrum and Manley, 2015; Quinn and Brannon, 1982; Stalleicken et al., 2005; von Frisch, 1949; Wiltschko et al., 1998b). Furthermore, cue hierarchies may be context-dependent, so that even in two situations where the available environmental cues are the same, an animal’s preferred cue may not be consistent. For example, a preference for light or dark in zebrafish *Danio rerio* depends upon both the surrounding light levels and the presence or absence of olfactory food stimuli (Stephenson et al., 2011). In other words, fish favour darker, presumed safer environments in the absence of food, but seek lighter ones when they can sense prey. Such flexibility of behaviour can manifest in animals as a consequence of learning about cues and their role for tuning other sensory systems. Additionally, behavioural changes may occur as animals enter new life phases where other habitat requirements become relevant ontogenetically (Brant et al., 2016; Dixson et al., 2011; Forward et al., 2003; Hawryshyn et al., 1990; Liu et al., 2016; Minot, 1988; Timmermann and Plath, 2009). A prominent example includes the use of compass systems by some bats and birds; these systems are used at night but are calibrated by celestial cues around sunset (Able and Able, 1990; Buchler and Childs, 1982; Lindecke et al., 2019; Muheim et al., 2006). Temporal shifts in cue-specific responses can occur in reaction to cue availability fluctuations or in accordance with circadian and circannual rhythms. For instance, cloud cover during migration season can disrupt the use of celestial cues and induce migratory birds to modify their flight trajectories and orientation mechanisms, switch to alternative cues for navigation, or even postpone their migration (Emlen, 1967; Griffin and Goldsmith, 1955). Therefore, when investigating an animal’s behavioural response to a sensory stimulus, it is paramount to consider that even when cues are consistently present in an (experimentally controlled) environment, the responses to these cues can fluctuate over different temporal scales aligned with environmental/biological rhythms or ontogeny. A limited understanding of this element of context may lead one to believe that a particular cue is unimportant when, in fact, it may not have been presented to the animal at the time when it is specifically relevant.

Here we investigate whether migratory bats have a temporally varying preference for light over the course of an 18-hour period. This timeframe spans from the usual onset of activity in the evening within a bat roost, through their nocturnal flight activity, to the early morning hours when migratory bats occupy stopover roosts (Erkert, 1982). Perhaps surprisingly, moving towards (artificial) light in night-active animals has been shown in many species including insects (Owens and Lewis, 2018), birds, and sea-turtles (McLaren et al., 2018; Witherington, 1997), as well as in bats (Chase, 1983; Lindecke et al., 2021; Voigt et al., 2017). This seemingly paradoxical light preference raises questions about the functional significance of nightly light-seeking behaviour. An attraction to artificial light at night (ALAN) is generally considered to be harmful for animals (Davies et al., 2014); causing unnatural behaviours (Botha et al., 2017), increased predation (Nuñez et al., 2021), and population declines (Sanders and Gaston, 2018). In the case of bats, responses to ALAN vary across species, often depending on their foraging ecology (Voigt et al., 2021). Under ALAN, free-flying bats are put off from drinking, but conversely continue (and some increase) foraging behaviour, exploiting congregations of insects under street lighting (Mathews et al., 2015; Russo et al., 2019; Schoeman, 2016). Negative effects of ALAN on bats can cause unhealthy bat populations (Boldogh et al., 2007); eventually, bats may leave habitats where ALAN is present (Rydell et al., 2017; Stone et al., 2009). ALAN can act as a sensory trap by causing bats to reduce call emissions in lit environments, resulting in collisions that could be avoided if using echolocation (Bradbury and Nottebohm, 1969; Orbach and Fenton, 2010; Test, 1967). Yet, despite the dangers potentially presented by light, several species preferred lit exits in Y-maze tests in which they had to choose between lit and dark routes (Chase, 1981; Chase, 1983). This preference has been found to modulate with time-of-day in species known as “cave-dwelling” and is at its highest when bats begin to leave their large roosts (Chase, 1981). It is thought that this circadian modulation to orient towards lit environments is explained by the animals’ use of the last daylight entering the cave entrance as a navigational cue to guide their exit when it is time to forage.

We have applied the methodology of (Chase, 1981) in conducting Y-maze choice experiments, except that our study focuses on a long-distance migratory species, the Nathusius’ pipistrelle bat, *Pipistrellus nathusii*. This species is not typically cave-dwelling but mainly roosts in trees (tree-dwelling bats), using crevices and cavities, both in summer and also often in winter (Gottfried et al., 2019). *P. nathusii* tolerate or even orient towards light during walking locomotion in Y-mazes (Lindecke et al., 2021) and in free flight during migration (Voigt et al., 2017; Voigt et al., 2018). This preference for light is not easily explained by a need for a cue to navigate towards a cave or roost exit that is beyond the echolocation range, as it is in cave-dwelling bats. Rather, we hypothesise that light-seeking behaviour in *P. nathusii* may be linked to their long-distance migratory lifestyle and that their photic orientation is subject to a circadian modulation reflecting seasonally-specific needs. For example, it is known that migratory soprano pipistrelles, *P. pygmaeus* which are also tree-roosting bats, use the solar disc as a directional cue for magnetic compass calibration during sunset (Lindecke et al., 2019; Schneider et al., 2023). Hence, the function of light cues in the behaviour of tree-dwelling bats, such as *Pipistrellus* species, may be connected in part to the mechanisms underlying migratory navigation.

Our fundamental premise of context-dependent light/dark preferences in *P. nathusii* rejects the notion of a rigid, continuous preference for the lit exit, i.e., phototaxis. Instead, we predict that the importance of light as a cue for these bats in a dark, roost-like environment peaks around sunset, driven by two non-mutually exclusive mechanisms: at this time, migratory pipistrelles may either use the Sun’s azimuth to determine their migratory direction (‘compass-calibration hypothesis’) or leave their stopover roosts to forage on insects that are active before a nightly decrease in temperature reduces their availability (‘refuelling hypothesis’). We also expect the light preference to reverse at least shortly before sunrise, as migrants need to seek roosts to recover from migratory flight (Costantini et al., 2019), and therefore would orient towards dark crevices to be protected also from predators during their daytime rest at stopover sites (‘roost-search hypothesis’) (Fig. 1). In contrast, under the refuelling hypothesis, bats may continue foraging as insect activity increases again under rising light levels and temperature at dawn (Learner et al., 1990; Reynolds et al., 2008). However, the ‘roost-search’ hypothesis, here in the context of migration, is closely connected also to the ‘predation’ hypothesis (Duvergé et al., 2000; Jones and Rydell, 1994), which aims to explain bat roost-emergence and re-entrance as a function of predator avoidance; bats would emerge as late as possible and return as early as possible to avoid predation. However, if they miss sunset compass-calibration or have heightened energy demands after long migratory transits, bats could again favour the lit exit at dawn to attempt foraging before roosting. This contrast between the roost-search and refuelling hypotheses, as well as compass-calibration, at dawn provides a valuable framework for understanding the relative importance of light cues in *P. nathusii*’s daily cycle.

**Fig. 1.**
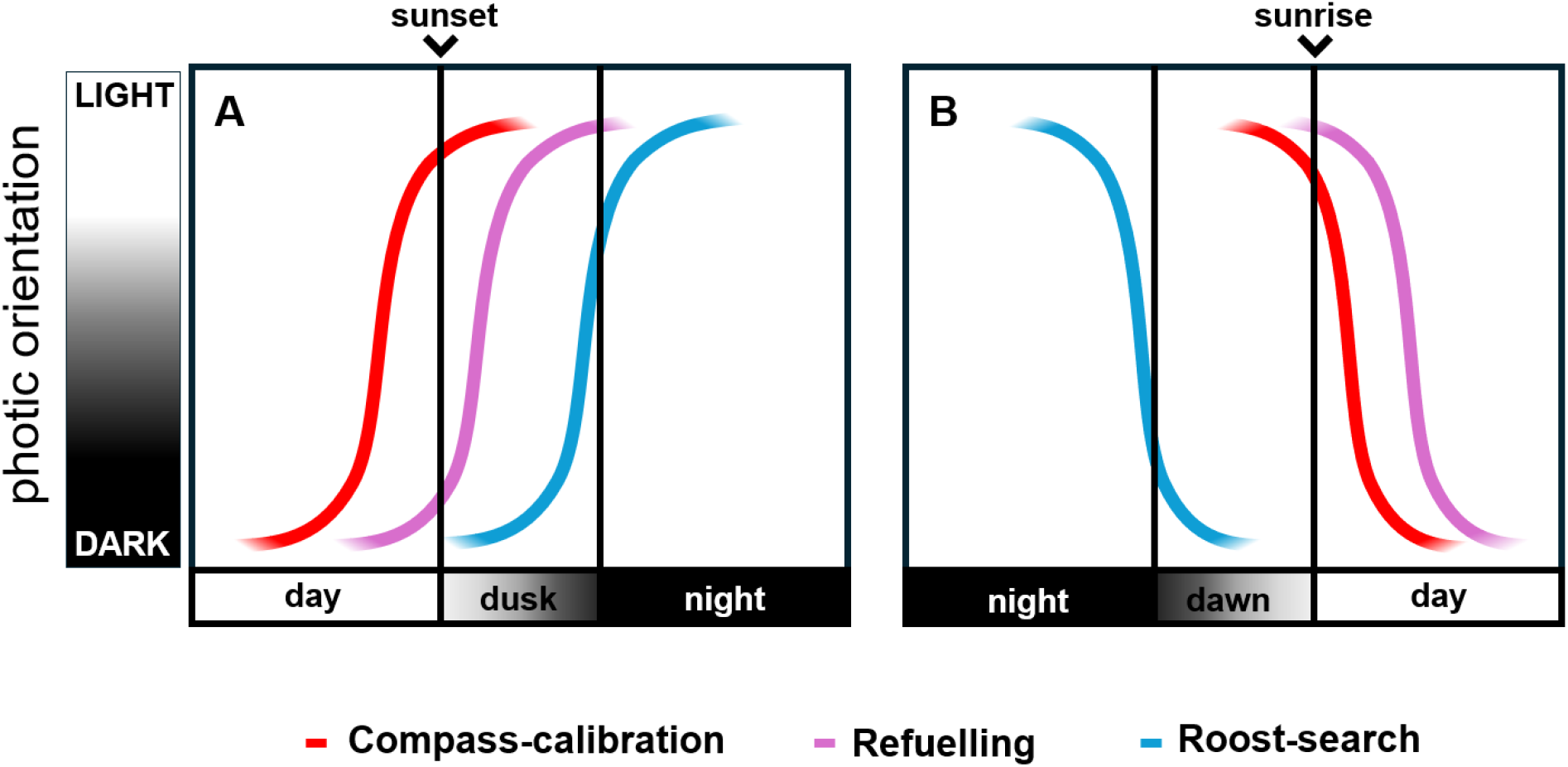
Schematic illustration of theoretical dynamics of light/dark y-maze exit preference (coloured lines) as a function of temporal light changes. The y-axis indicates the tendency of bats to exit via the lit or dark arm. On the x axis, day becomes dusk when the sun drops below the horizon, and dusk becomes night when the sun has dropped 18 degrees below the horizon, at which point there is a total absence of sunlight. This process is reversed for sunrise. At the onset of the bat activity period (A), under the ‘compass-calibration’ hypothesis, we predict bats to leave prior to sunset to get a view of the solar disc setting location (A, red line). According to the ‘refuelling’ hypothesis, bats leave after sunset during dusk to take advantage of a peak in insect abundance before nightfall (A, violet line). For the ‘roost-search’ hypothesis, bats should begin to select the lit arm only at the end of the dusk period to avoid predation (A, blue). Prior to daybreak (B), if bats use the rising sun in their compass calibration they will begin to choose the dark arm around sunrise (B, red line). In this scenario, the rising sun provides the necessary cue for orientation. For migratory bats needing to refuel, a preference for light should last beyond sunrise to capitalise on prey availability (B, violet line). Roost-searching bats should switch to dark preference before dawn (B, blue line) in order to find a dark roost before predation risk increases.

The impacts of this research are threefold: firstly, it will clarify cue hierarchies in a migratory bat. Secondly, it will inform how and when experiments involving light cues should be conducted in *P. nathusii* and comparable species of bats. This is vital for current researchers who seek to establish the sensory cue-use and hierarchy that informs migratory bat navigation (Lindecke et al., 2015; Lindecke et al., 2021; Schneider et al., 2023). Thirdly, a comprehensive understanding of nocturnal animals’ attraction to light is essential for evidence-based environmental planning and wildlife-friendly lighting design (Laforge et al., 2019; Stone et al., 2015).

## MATERIALS AND METHODS

### Experimental setup

We used a Y-maze constructed from dark brown film-faced plywood (Lindecke et al. 2021); the surface colour minimized light reflections. A 100 mm long runway with a cross-section measuring 80 mm wide and 60 mm heigh led to the ‘arms’ which were each 200 mm in length. The maze was inclined towards the exits 10° following recommendations of previous work (Chase, 1981). The floor was lined with a textured, polyvinyl chloride (PVC) material that ensured good grip for crawling bats and effective cleaning to eliminate scent trails that might influence bat decision-making. Light levels were measured using an LX-1102 light meter (0.01 lx resolution, ± 3% rdg.; Lutron Electronic, Taiwan). The exit of one arm was illuminated at 120 lx using dim white light (Ledlenser GmbH & Co. KG, Solingen, GER), which was reflected off a brown hardboard wall 1 m away. This setup ensured that the bats could not use echolocation to detect the three LEDs that provided the light source. The lighting condition resembles the dim light regime after sunset on a cloudy day, where the light levels can range between 100 to 150 lx, depending on cloud cover (Hölker et al., 2021). At the same time, this light level approximates the diffuse illumination under a canopy during the two hours before sunset (Hölker et al., 2021). However, considering the roosting ecology of *P. nathusii*, any brighter conditions would have been unrealistic for a simulation of a roost-emergence, as their roosts are generally located within or below tree canopies or near forest edges, typically lacking a clear view of the horizon (Flaquer et al. 2005; Gelhaus and Zahn 2010). A light-blocking barrier between the arms of the maze ensured that the other exit remained comparatively dark, with a light level of 0.01 lx, which resembles a star-lit night during a new moon (Kyba et al. 2017). The area where the arms of the Y-maze diverged (choice point for a bat) received indirect illumination of 0.12 lx from the lit arm, which is similar to the conditions at the choice point in previous studies in insectivorous yet cave-dwelling bats (*Tadarida brasiliensis*; Mistry & McCracken 1990, Mistry 1990). At the entrance of the maze the light level was 0.02 lx. Other than the maze lighting, the testing room was kept dark. Bats were tested in alternate order, with the lit arm of the maze changing for each trial.

### Experimental bats and site

Adult *Pipistrellus nathusii* (Keyserling & Blasius, 1839) were caught at Pape Ornithological Station (University of Latvia, 56°09’ N 21°03’ E, Kurzeme Municipality, Latvia) between 10 and 19 August 2018, using a funnel trap of the “Rybatschi type” (Keišs et al., 2021) as part of the long-term bat ringing programme. Bats were captured and handled under licence (Nr. 31/2018) from the Latvian Nature Conservation Agency adhering to the ASAB/ABS guidelines for the treatment of animals in behavioural research (ASAB, 2018). After controlling for seasonally appropriate body mass (≥7.0 g), bats were kept in wooden boxes in groups of 3-5 individuals in a dark and quiet keeping room until the following day. For the trials, all boxes were transferred to a separate building. During transfer, ventilation holes were covered to prevent exposure to daylight. The boxes were then placed in a dark room adjacent to the test room. We chose not to provide bats access to environmental light cues at any point before the trials to avoid biasing their photic orientation. This included preventing bats from seeing the previous sunset as a measure of experimental standardisation. Using an ultrasound bat detector (Echometer EM3 +, Wildlife Acoustics, Inc., Maynard, MA, USA), we confirmed that no bat calls were audible in the test room (from bats other than the one being tested). In total, we tested 304 bats, all of which were ringed upon release using metal ‘bat bands’ registered at the Latvian Ringing Centre. Individuals were tested only once and freed in the nearby coastal forest within at least 1 hour after a trial. Any bat tested after 4:00 AM, i.e., approx. 2 hours before sunrise, was kept for the day again, fed mealworms (*Tenebrio molitor*), given water, and released the following evening.

### Testing procedure

Trials were performed at room temperature. Each bat was carefully transferred by hand to the acclimatisation compartment of the Y-maze. After a 20-second period, a sliding door was opened so that the bat was free to crawl along the runway and reach the bifurcation. An exit choice was recorded when the bat emerged from one of the open arms of the Y-maze. We timed emergence latency, i.e. the time it took an animal to reach the exit. We monitored the bat activity and exploration behaviour through their vocalisations using an ultrasound detector (Magenta Bat5 Digital Quartz Bat Detector, Magenta Electronics Ltd., UK) with headphones tuned to 40 kHz and placed over the Y-maze. If a bat had not attempted to emerge from the Y-maze within 3 min, the trial was cancelled. This cut-off-time is derived from the experience of previous work in the same species (Lindecke et al., 2021). Clean sheets soaked with ethanol (70%) were used to remove any residue on the runways between trials.

Experiments were conducted between the 10th and 20th of August. During this period, the sunset/sunrise times shifted from 21:25/05:55 to 21:01/06:14. The mean night length was 9 hours 4 minutes, whilst the mean sunset/sunrise times were 21:10/06:06. Of the 304 bats tested, six did not emerge from the acclimatisation compartment and were therefore excluded from the analysis.

### Statistical methods

Light preference over time was modelled using a second order polynomial generalised linear model with a binomial distribution. Order of the polynomial was incremented until the best model fit was obtained. Two models were created each with time anchored on different events; model 1) time anchored to sunset, model 2) time anchored to sunrise. Latency to exit through time was modelled using a second order polynomial glm with a Gamma distribution. Emergence latency was compared using the Mann-Whitney U test since data were not normally distributed (P<0.05). All statistics were performed in RStudio Build 372 and R version 4.1.2.

## RESULTS

Bats always emitted echolocation calls when exploring the Y-maze. The probability of bats choosing the lit arm of the Y-maze varied significantly through time in a quadratic fashion (p = 0.001, z = -3.18, Fig. 2). The chance of a bat preferentially choosing the lit exit first rose above 0.5 at 2 hours 21 mins before sunset, this increased to a peak of 0.67 (95% CI: 0.58 to 0.74) at 2 hours 41 mins after sunset (Fig. 2A). The chance of light-selection then fell below 0.5 at 1 hour 15 mins before sunrise and by sunrise this had reduced to 0.40 (95% CI: 0.30 to 0.52) Fig. 2B). We found no differences between sexes (p = 0.578, z = -0.56), no side preference (p = 0.670, z = 0.390), and no changes throughout the migratory period (p = 0.542, z = 0.609) in light/dark selection. Latency to choose an exit did not change throughout the night (p = 0.193, t = 0.218), nor did it differ between sexes (p = 0.703, t = -0.382) or between the bats that chose light or dark exits (p = 0.215, t = -1.243).

**Fig. 2.**
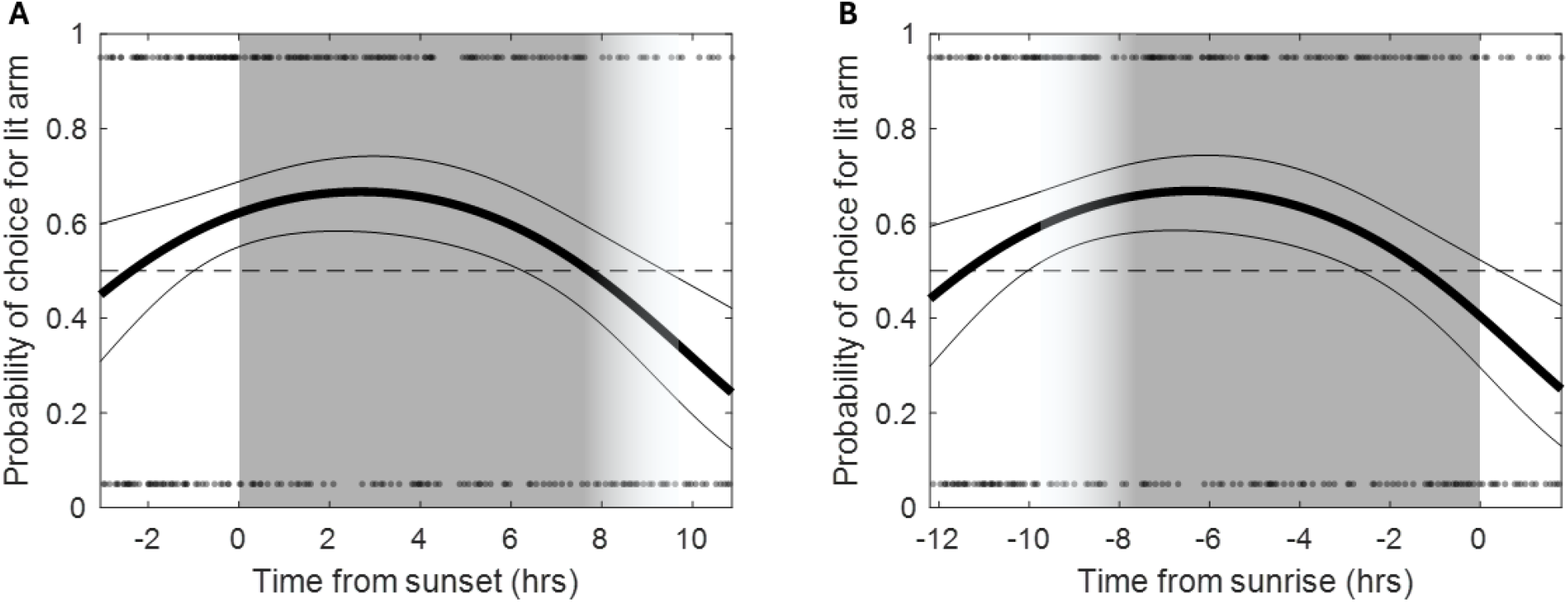
Timing of circadian shifts in light/dark preference. The probability of bats to choose the lit arm of the Y-maze is shown in both A and B with a thick black line. The upper and lower 95% confidence intervals are shown with thin black lines either side. In A, model predictions used the time-from-sunset in hours, whilst in B the time-from-sunrise was used. Small circles show the light/dark choices of all individuals at the top/bottom of both graphs respectively.

## DISCUSSION

We found pronounced circadian shifts in the light-dependent decision-making of migratory *P. nathusii* released from a Y-maze. While the bats generally preferred the lit exit for most of the night, the strength of this preference changed throughout. Preference for the lit exit rose before sunset and shifted to the dark exit towards the end of the night. These findings suggest that *P. nathusii* use light cues to guide exit-route choice, implying a more complex decision-making process for emergence than a simple response to light intensity.

### Support for the compass calibration hypothesis

Migratory soprano pipistrelles (*P. pygmaeus*) have been shown to calibrate a magnetic compass at sunset using the Sun’s position (Lindecke et al., 2019; Schneider et al., 2023). This behaviour could be a conserved trait among migratory bats, or at least, among pipistrelle species. Our observation that *P. nathusii* increasingly chose the lit exit in the two hours before sunset supports the possibility that light cues could be integral to compass calibration in this species as well. Notably, for a free-flying *P. nathusii* at our study site, the Sun’s disc would have been visible at a maximum elevation of 18° above the horizon at this time of the day (Hoffmann, 2025). This assumes unobstructed viewing and clear weather conditions. In the wild, the light level we used could indicate that the Sun is setting outside, prompting *P. nathusii* to emerge and access solar cues in open space more directly.

The persistence of orientation toward the lit exit well beyond the sunset period suggests that *P. nathusii* may opportunistically use available light cues for orientation. Importantly, the period when the solar disc is still visible at the horizon may provide animals with a uniquely precise reference point to calibrate their (magnetic) compass system for later use. Once the Sun descends below the horizon, this possibility vanishes: the remaining glow is too broad and diffuse to offer the same accuracy to anchor such calibration. However, the post-sunset glow may still serve bats as a landmark for immediate orientation until the end of dusk. Still, such landmark-based use of light differs fundamentally from compass calibration, as it only works while the cue is directly perceived. Consistent with this, Buchler and Childs (1982) demonstrated that big brown bats could use the post-sunset glow to head into their colony-specific direction of foraging grounds, but after dusk, without access to the glow, they departed in random directions. This strongly suggests that post-sunset glow does not calibrate a magnetic compass but rather acts as a temporary directional landmark.

Comparable behaviours have been reported in night-migratory birds. Many species take off for transit flights within a few hours of sunset likely using the post-sunset glow for orientation (Moore, 1986; Muheim et al., 2006). Some songbirds could further be shown to recalibrate their compass not only at sunset, but also at sunrise, using the polarized light pattern in the sky at twilight (Muheim et al., 2006; Muheim et al., 2007; Muheim et al., 2009). Yet, a previous study in migratory *P. nathusii* at our field site showed that polarized skylight during sunset was not effective for calibrating their compass system (Lindecke et al., 2015). Our observation of bats’ preference for darkness during the early morning hours indicates that sunrise is not typically used for recalibration in this species, unlike in certain migratory birds. Accordingly, the bats’ preference for the dark exit of the Y-maze during the early morning hours may be better explained by the roost-search hypothesis.

### Support for the roost-search hypothesis

The counterintuitive preference for light at night aligns with previous studies on visually guided emergence behaviour in cave-dwelling bat species (Chase, 1981; Gorresen et al., 2015; Mistry and McCracken, 1990). However, a key difference is that during daytime, *P. nathusii* in our study preferred the dark arm of the Y-maze, whereas cave-dwelling species tested at similar times kept a preference for the lit arm (Chase, 1981; Mistry, 1990). In species like Seba’s short-tailed bat (*Carollia perspicillata*) or Mexican free-tailed bat (*T. brasiliensis*), orientation toward darkness matches the timing of return to caves at sunrise. Surprisingly, this preference reversed to a preference for light within two hours after sunrise (Chase, 1981; Mistry, 1990). By contrast, we observed no such change in *P. nathusii*, which is consistent with an urgency to locate and remain in a secure roost, avoiding brighter and exposed locations during daytime. Notably, neotropical Pallas’s mastiff bats (*Molossus molossus*) also chose the dark arm for part of the night when they paused foraging to seek temporary refuge in their roost (Chase, 1981). At our site, *P. nathusii* flight activity typically does not decline mid-night under favourable weather (Šuba et al., 2012). Hence, we would not expect a temporary shift of preference for darkness in the Y-maze, like that seen in *M. molossus*, but rather a continued preference for the lit exit throughout most of the night, with a switch toward exiting into the dark only as sunrise approaches. Together, these patterns suggest that choosing a dark route in a Y-maze generally indicates roost-search behaviour in bats.

### Little support for the refuelling hypothesis

Finally, we considered whether light preference could be linked to foraging motivation. The highest preference for light near sunset and in the early part of the night broadly aligns with the time when *P. nathusii* typically leave their roosts and forage during the non-migratory season (Ciechanowski et al., 2009). However, in our study, bats exited the illuminated arm significantly earlier than expected if timed solely for peak insect availability after sunset. Notably, acoustic monitoring at our site provides no evidence of bat foraging activity before sunset (Šuba et al., 2012), suggesting that, at least during migration, pre-sunset orientation toward light in the Y-maze is not explained by foraging motivation. At sunrise, insect activity usually increases with rising light levels and temperature (Learner et al., 1990; Reynolds et al., 2008), but rather than continuing to choose the lit exit to exploit dawn conditions for foraging, *P. nathusii* in our study switched to the exit-route leading to darkness. This behaviour suggests that roost-search and predator avoidance commonly override other needs, i.e., refuelling.

### Spatial cognition, cue-hierarchy and decision making

Our findings highlight a cognitive dimension of the bats’ behaviour: light/dark choices in the Y-maze can be interpreted as context-dependent weighting of sensory cues. Rather than reflecting a rigid phototaxis, the bats’ responses appear gated by circadian state, consistent with state-dependent decision thresholds. In this view, light functions as a cue whose utility varies with time of day and migratory priorities, flexibly re-weighted against other sensory inputs such as echolocation. This pattern exemplifies temporal gating, where the relevance of a cue is not constant but modulated by circadian phase, effectively switching the behavioural outcome of the same stimulus (Brown, 2016). Within this framework, we propose a ‘light-mediated space association’ hypothesis: bats cognitively map illumination onto functional spatial categories, with light signifying open, exterior environments and darkness signifying enclosed, roost-like conditions. Such associations help to explain why many bat species, including *P. nathusii*, orient towards lit exits around sunset but switch to dark exits before sunrise, when roost-search and predator avoidance are critical (Chase 1981; Mistry 1990; Lindecke et al. 2021). Unlike cave-dwelling species, whose choice of a lit exit may reflect disturbance-induced escape when they were tested inside their colonie’s familiar roost (Mistry & McCracken 1990), migratory *P. nathusii* in our study likely used light as a proxy for ‘outside’ under controlled yet ecologically relevant conditions. This interpretation integrates our results with earlier findings and generates testable predictions: if light preference reflects cue-based space categorisation, then bats from dark-sky versus light-polluted populations, as well as naïve subadults with little prior exposure to artificial light, should differ in the strength and timing of their light-dependent orientation. Such comparative approaches would help determine whether nightly light-seeking in *P. nathusii* is primarily a migratory adaptation or a response shaped by anthropogenic environments.

Modern, anthropogenically lit landscapes can blur a natural association of light and space conditions. Preferences for a lit exit by city-dwelling bat species after sunset could indicate an experience-based association of illuminated conditions with “outside” environments in light-polluted areas where natural twilight is absent (Barré et al. 2022). This behaviour may be beneficial for exploiting concentrated insect prey near artificial lights. Indeed, *P. nathusii* are not necessarily repelled by artificial light sources and are known to exploit illuminated environments, such as urban areas or other light-polluted habitats, where insect prey can gather in dense concentrations (Lewanzik and Voigt 2017; Voigt et al. 2017). While such behavioural adaptation highlights the species’ resilience and ability to exploit anthropogenic changes in their environment, this lack of light-aversion also raises conservation concerns as artificially lit areas might expose bats to higher predation risk or disrupt natural migratory timings by masking circadian cues (Smith et al., 2021).

### Limitations of the y-maze approach

While the Y-maze design allowed us to isolate and quantify the influence of light under controlled conditions, it inevitably restricts the full spectrum of natural stimuli bats may integrate in the wild, such as auditory, thermal, or olfactory signals. Although these design choices helped us isolate the role of light in exit-route decision making, future experiments could incorporate naturalistic lighting transitions or limited feeding opportunities to mimic field conditions more closely. Moreover, quantifying bat calls could clarify whether vocal activity varies with time of day or exit choice. Our findings, together with recent laboratory work showing that echolocation performance in *Myotis daubentonii* is stereotyped even in lit conditions (Uebel et al., 2024), suggest that echolocation is largely reflex-like, whereas vision provides the decisive cue for higher-level exit-route choices in migratory bats. Echolocation may provide complementary information to vision, particularly for accurate distance measurement, as suggested after observations of bats calling regularly even in bright daylight (Eitan et al., 2022; McGowan and Kloepper, 2020).

### Conclusions

Our findings are evidence of a nuanced, time-dependent reliance on light cues by migratory *P. nathusii* in a Y-maze. The bats shifted between light and dark preference in accordance with multiple ecological and navigational requirements, including compass calibration, potential refuelling, and timely roost-search. This time-dependent shift in orientation challenges the idea of simple, continuous phototaxis in nocturnal bats. Although echolocation is crucial for short-range manoeuvring, vision evidently governs higher-level spatial decisions, underscoring a multimodal sensory strategy in migratory bats. Further, it is suggested that bats may associate light with open-air foraging spaces and darkness with roost-like environments (*light-mediated functional space association hypothesis*). Nightly light-seeking, also observed in other bat species, may reflect an evolutionary remnant of diurnally active ancestry. If so, the association of light with open-air spaces would not only be a context-dependent decision but also a vestigial trait modulated by circadian state.

Beyond its relevance to *P. nathusii*, this study offers broader implications for laboratory protocols that examine bat sensory hierarchies or cue-conflict scenarios (e.g., in magnetoreception research or social behaviour). Researchers must ensure that experiments align with bats’ natural circadian timing of cue use; otherwise, critical behaviors may go undetected or be misinterpreted. Finally, the interplay between circadian rhythms and anthropogenic light has direct conservation implications. Expanding light pollution threatens to disrupt bats’ precise balance of risk avoidance (roost search), compass calibration, and refuelling. Wildlife-friendly lighting strategies, informed by species-specific sensory responses, will be critical in minimising adverse impacts on migrating bats and other nocturnal organisms.

## Acknowledgment

We thank the Pape bat catching team, esp. Ilze Kukare, Normunds Kukare and Roberts Jansons for their help in the field. We are grateful to Christian C. Voigt and Oskars Keišs for logistical support of this project.

## Author contributions

Conceptualization: O.L.; Methodology: O.L.; Investigation: D.V., O.L., V.V.; Data curation: O.L., D.V., W.T.S.; Formal analysis: W.T.S., O.L.; Funding acquisition: O.L.; Resources: O.L., D.V., V.V.; Supervision: O.L.; Visualization: W.T.S., O.L.; Writing – original draft: O.L., W.T.S.; Writing – review & editing: O.L., D.V., V.V., W.T.S.

